# A Model for Self-Organization of Sensorimotor Function: The Spinal Monosynaptic Loop

**DOI:** 10.1101/2021.05.18.444712

**Authors:** Jonas M.D. Enander, Adam M. Jones, Matthieu Kirkland, Jordan Hurless, Henrik Jörntell, Gerald E. Loeb

## Abstract

Recent spinal cord literature abounds with descriptions of genetic preprogramming and the molecular control of circuit formation. In this paper we explore to what extent circuit formation based on learning rather than preprogramming could explain some prominent aspects of the spinal cord connectivity patterns observed in animals. To test this we developed an artificial organism with a basic musculoskeletal system and proprioceptive sensors, connected to a neural network. We adjusted the initially randomized gains in the neural network according to a Hebbian plasticity rule while exercising the model system with spontaneous muscle activity patterns similar to those observed during early fetal development. The resulting connection matrices support functional self-organization of the mature pattern of Ia to motoneuron connectivity in the spinal circuitry. More coordinated muscle activity patterns such as observed later during neonatal locomotion impaired projection selectivity. These findings imply a generic functionality of a musculoskeletal system to imprint important aspects of its mechanical dynamics onto a neural network, without specific preprogramming other than setting a critical period for the formation and maturation of this general pattern of connectivity. Such functionality would facilitate the successful evolution of new species with altered musculoskeletal anatomy and it may help to explain patterns of connectivity and associated reflexes that appear during abnormal development.

## Introduction

Motor commands from the brain are processed through the neuronal circuitry of the spinal cord, where they are integrated with sensory signals before producing output in Sherrington’s final common pathway of the motoneuron (MN) (Sherrington, 1906). Because the sensory signals depend on the musculoskeletal mechanics (the ‘plant’ in engineering terms), the combination of spinal circuitry plus plant mechanics defines the control problems that the brain must solve to generate desired motor behaviors (Loeb et al., 1990, 1999, 2011). It would seem to be advantageous for the spinal circuitry to reflect the plant mechanics (Nichols, 1994), but it isn’t clear what that means given the complexity of both the mechanics and the circuitry in vertebrates.

The recent spinal cord literature abounds with descriptions of genetic preprogramming and the molecular control of circuit formation (Delile et al., 2019; Hoang et al., 2018; Lai et al., 2016; Osseward and Pfaff, 2019). The question we are asking is whether the genetic programs specify only general rules for types of connectivity, rather than being responsible for creating fully functional connectivity patterns that reflect the functional properties of the musculoskeletal plants. One example of such a general rule for connectivity formation would be the genetic definition of a subset of spinal interneurons as commissural interneurons, which allows them to grow their axons across the midline, whereas other spinal interneurons and sensory afferents do not (Lanuza et al., 2004). Such ontogenetic specification could, for example, also be important for the topological partitioning of the spinal termination zones (Sürmeli et al., 2011; Zlatic et al., 2009). We here explore to what extent circuit formation based on learning rather than preprogramming could explain some functional aspects of the spinal cord connectivity patterns observed in adult animals.

Fetal animals generate spontaneous motor twitches that would provide coherences between sensory and motor signals that reflect the mechanics of the musculoskeletal system and could be used for aspects of self-organization (Mendelson and Frank, 1991), for example pruning of incorrect connectivity and specification of synaptic weights. Specifically, it is well known that the spindle primary afferents (Ia) project monosynaptically and predominantly to the homonymous αMNs (alpha, extrafusal) that innervate the extrafusal muscle fibers in the muscle in which the spindle is located, giving rise to the classical stretch reflex. However, the Ia to αMN connectivity is not confined to homonymous αMNs; Ia afferents can also make monosynaptic connections with the αMNs of other muscles, particularly those that have a related function (Eccles et al., 1957; Fritz et al., 1989; Hongo et al., 1984). That observation would suggest that learning, perhaps even before birth, can play a role in establishing this connectivity.

Activation of the homonymous αMNs will tend to shorten the muscle and its spindle receptors, reducing and even silencing spindle afferent activity. A Hebbian learning rule based on enhancing synapses in which there are simultaneous pre-and post-synaptic firing (“neurons that fire together wire together”) would then eliminate rather than promote homonymous Ia-MN connectivity. This type of learned connectivity has previously been achieved by employing reversed Hebbian learning (Petersson et al., 2003), while interesting it is not in line with what is known about spinal learning or typical long-term potentiation.

In mature mammals, spindle afferent activity can also be driven by an independently controlled γMN (gamma, fusimotor) system that causes contractions or stiffening of various intrafusal muscle fibers and thereby increases the excitability of both the primary (Ia) and secondary (II) sensory endings of the muscle spindle (Loeb, 1984; Loeb and Hoffer, 1985; Prochazka et al., 1988). However, γMNs differentiate and modulate spindles postnatally and later than αMNs and βMNs (Shneider et al., 2009). It is often forgotten that a substantial percentage of the spinal MNs in all vertebrates (including mammals) are, in fact, βMNs, which innervate both intrafusal and extrafusal muscle fibers (reviewed by Manuel & Zytnicki (2011)). We hypothesize that the widespread fusimotor effects of βMNs will be sufficient to overcome the relatively weak extrafusal contractions during early development, resulting in firing patterns in spindle primary afferents and MNs that could support Hebbian organization of the spinal monosynaptic reflex between them. Furthermore, this would be even more plausible in light of the presence of short duration muscle twitches known to exist early in development (Corner, 1977), which would generate sensory feedback while avoiding the spindle unloading caused by skeletal motion and muscle shortening.

In order to test the feasibility of self-organization of sensorimotor function based on musculoskeletal mechanics and behavioral experience, we have designed a model creature with the simplest musculoskeletal system that we thought might exhibit an interesting range of behaviors. Therefore, the model system does not feature a full replication of the macroscopic configuration of an extant biological musculoskeletal plant but is kept to a minimum with two arms and one flexor and extensor attached to each of the two arms. Indeed, skeletal linkages with higher number of degrees of freedom can become highly difficult to control, and an attempt to replicate them would risk consuming most of the energy on solving the high-dimensional control problem rather than the more fundamental issues we initially aimed to address. Our musculoskeletal plant is controlled by a spinal neuronal circuitry using a linear summation neuron model (LSM) (Rongala et al., 2021) which is a simplified version of a previously published implementation (Rongala et al., 2018) together with synaptic plasticity based on the Hebbian-inspired calcium co-variance learning rule (Sejnowski, 1977). In this paper, we describe how random twitching activation patterns, such as generated in the fetal spinal cord, could give rise to naturally occurring connectivity patterns of spindle Ia afferents to MNs, one of the first circuitry formation events during fetal development (Chen et al., 2003). These are the first steps toward identifying how the CNS might self-organize so as to learn how to get its musculoskeletal system to perform useful tasks.

## Methods & Model Design

The model system is depicted schematically in Figure 1, consisting of the organism (called an Oropod as explained below), a bounded World in which it can move, and one or more Objects with their own behavior with which the Oropod can interact (Fig. 1A–D). The Oropod is designed to be akin to a one-dimensional cephalopod, with two tentacle-like limbs that can be moved (up to ±4 units along the x-axis) so as to contact each other (Fig. 1B), to shift the Oropod’s position by pushing against the World boundary (Fig. 1C) and to contact and capture Objects (Fig. 1D). The Oropod can go through stages of sensorimotor development by first operating in isolation as described in this paper, then with added Objects with increasing complexity of their autonomous behaviors, and finally by changing its location in the World. The expected behavioral repertoire will be limited to that of submammalian vertebrates, whose anatomy and physiology can provide some useful hints for the design of the Oropod.

**Figure 1.**
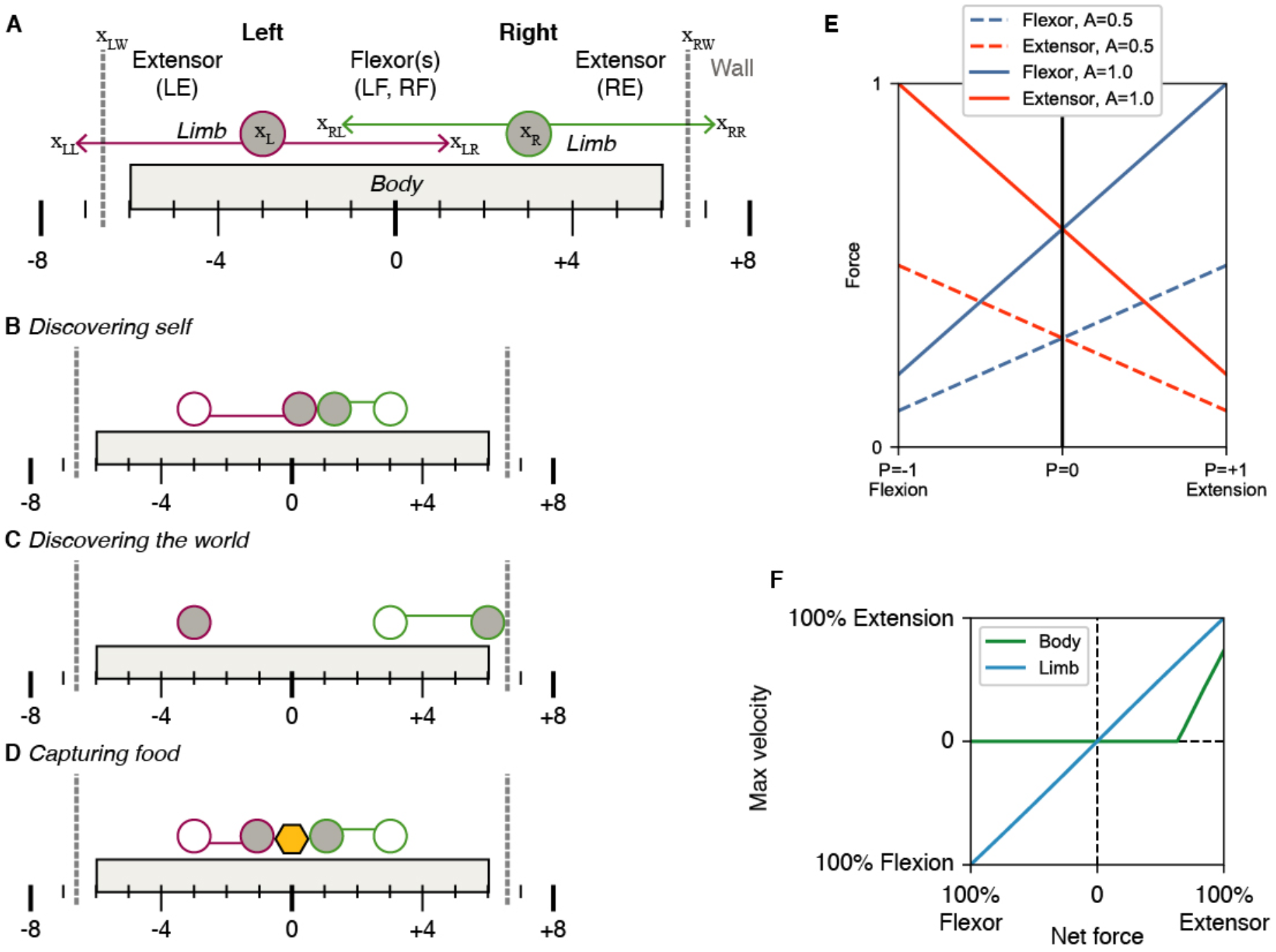
Macroscopic anatomy and behavior of the Oropod model system. A) The Oropod anatomy with Body, limbs and corresponding muscles Left Extensor (LE), Left Flexor (LF), Right Flexor (RF) and Right Extensor (RE). Left and right walls of the Oropod World are indicated with gray dashed lines, and the absolute global positions are indicated below the Body, and position terms used in equations 1–4 are pointed out with relative zero at x_LL_. The limbs’ anatomical limits are +- 4 units, as shown. Notable states of the Oropod are shown in panels B) ‘Discovering self’; C) ‘Discovering the world’ D) ‘Capturing food’ with the limbs. E) Graph of the force output (y-axis) relative the position of a limb (x-axis) and activation of the muscles (line colors according to legend). F) Graph over max velocity as a function of net force. Please note that due to stiction the body cannot move if the net force is not above the threshold.

### Musculoskeletal Mechanics

Each one-dimensional limb is operated by an antagonistic pair of muscles that are unidirectional force-generators (springs that can pull but not push). Extensor muscles pull the limbs outwards relative the center of the Oropod body and flexor muscles pull the limbs inwards and towards each other, resulting in a total of four muscles (Fig. 1A).

While large terrestrial organisms have mechanics dominated by inertia and Newton’s second law (force = mass x acceleration), small, aquatic organisms (from which vertebrates evolved) have limb and body mechanics dominated by viscosity, i.e. the force required to move increases with velocity (force = viscosity x velocity; equation 1 & 2). The Oropod limbs and body incorporate such viscous damping, allowing us to simplify the muscle model by omitting the force-velocity relationship present in terrestrial organisms. Thus, the velocity of the Oropod’s limb motion is proportional to net force (Fig. 1F; Net force is the difference in active force generated by the antagonist muscles pulling the limb in opposite directions plus any external force from contact with the end-effector). Furthermore, similar to the biological world, in the Oropod model system the force output of a muscle is a function of both its neural activation and its kinematic condition, which gives rise to “preflex” responses to perturbations (Brown and Loeb, 2000). The Oropod muscles incorporate the well-known “springlike” property of muscles operating on the ascending limb of their force-length curve (Fig. 1E). The spring curves have a mid-range overlap so that the organism can control “stiffness” by using different levels of co-contraction of antagonists.

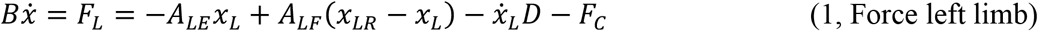

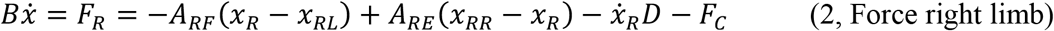

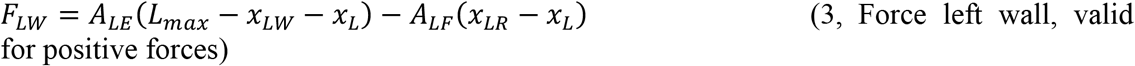

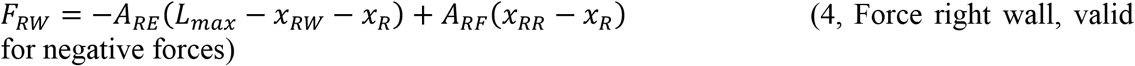

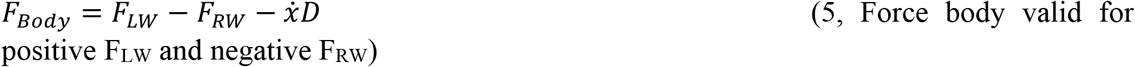

In the above equations describing the force relationships *B* denotes the viscosity, *x* is position relative most leftward possible limb position, *A* is muscle activation, *D* is damping due to viscosity and *L_max_* is maximal muscle length. Subscript L and R denote left and right respectively, and subscript LW and RW denote left and right wall respectively. Refer to Table 1 for a complete list of all variables used in the paper. Damping is a constant term due to our assumption that we model an organism dominated by viscosity and is set to 2.0 and thus renders the limb movements overdamped. Finally, F_C_ is external contextual forces which will be mostly due to contact with limbs, walls, prey and anatomical stops.

**Table 1.** Parameter definitions.

| Parameter | Symbol | Value or range |
| --- | --- | --- |
| Position along the x-axis relative a zero-point at the far left | x | -inf—inf |
| Damping | D | 2.0 |
| Muscle Length | L | 0—1 |
| Output from a neuronal compartment. Superscript plus denotes the positive part of the range. | A | -1—1 |
| Sign matrix denoting excitatory or inhibitory synapses | S | +1 or -1 |
| Synaptic weight | w | 0—1 |
| Learning signal | l | -1—1 |
| Kalman gain for Activity | K<sub>A</sub> | 0.3 |
| Kalman gain for Learning signal | K<sub>L</sub> | 0.001 |
| Kalman gain for Mean activity | K<sub>M</sub> | 4.0e-5 |
| Filter function for synaptic activity. High-pass (0.05 Hz threshold) for EPSP and low-pass (0.2 Hz threshold) for IPSP. | $\phi_L$ | Function |
| One-dimensional Kalman filter. | $\phi_K$ | Function |

The Oropod body also has stiction, represented as a threshold value of net force before the body starts to move, which avoids accidental drifts in location in its world as a result of cumulative effects of accelerating the small inertial masses that are required to avoid singularities in most physics engines. This stiction has been set fairly high (~50% of maximal muscle force, Fig. 1F) so that the Oropod maintains its current position even while pushing against the World with low forces.

As our aim was to model dynamics in the fetal stage when muscle strength is reduced compared to the adult stage (Gokhin et al., 2008), we had to define muscle strength as a feature that could change with maturation. Accordingly, we defined full muscle strength to be when the organism could move one limb from one extreme position (full flexion/extension) to the other in a short but not unreasonable time (Fig. 4A). This was defined as roughly 2 seconds from an animation of the model, and the muscle force at this point was defined as 100% muscle force. For the main simulations of the organism the muscle force was reduced to 10% to model the decreased strength during the fetal stage.

**Figure 2.**
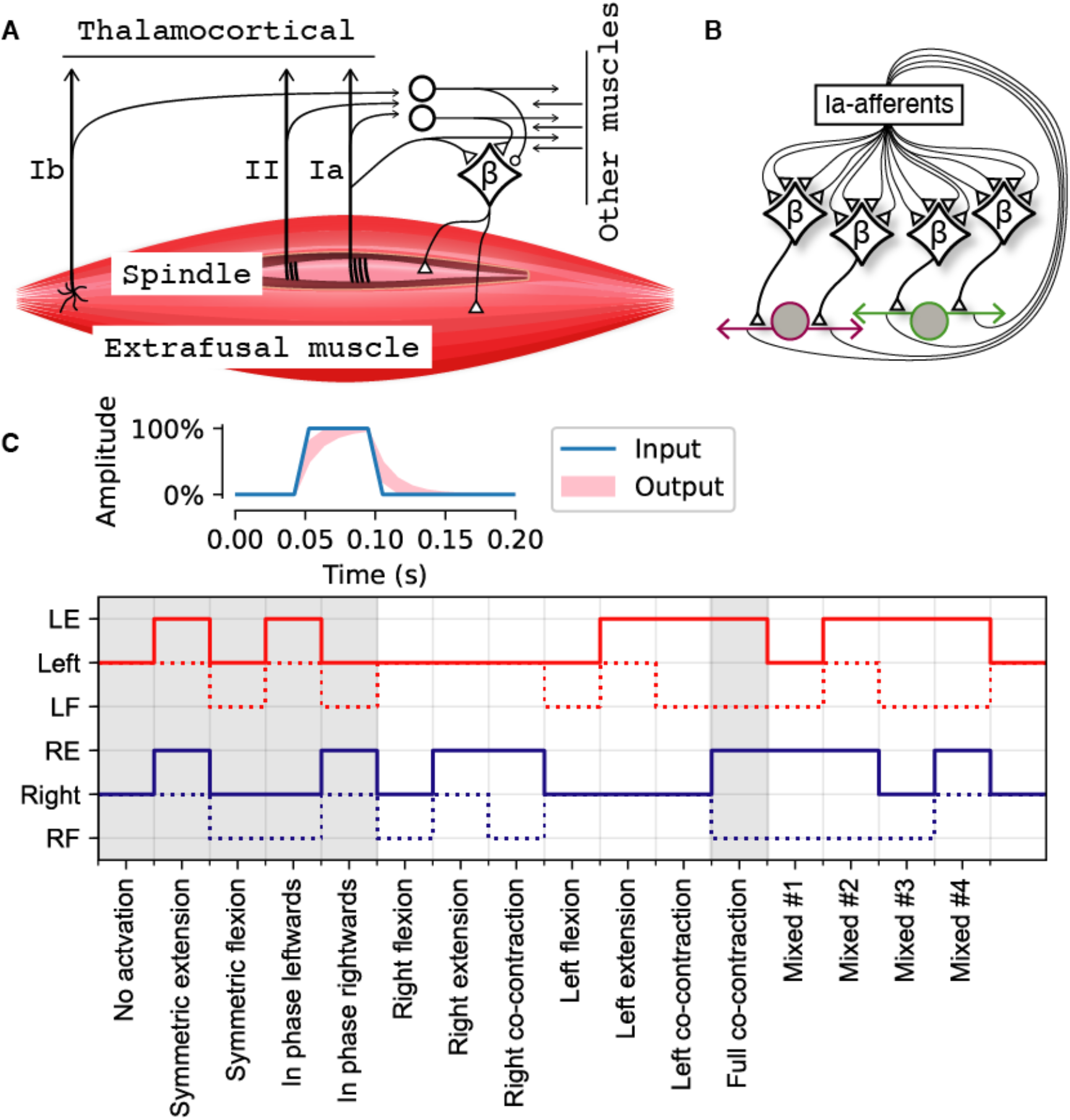
Muscle and neural network properties of the Oropod. A) Proprioceptive receptors of the Oropod muscle. B) Illustration of the neural network design of the present version of the Oropod. From each muscle Ia afferents synapse onto βMNs that project back unto their respective muscle (both extra-and intrafusal muscle fibers). Limbs and muscles are drawn as in Fig. 1 A–D. C) In the top is an example APG activation with 100% amplitude and 50 ms duration, the blue line shows the internal square pulse into the APG and the light red area shows the output range when fed through the one-dimensional Kalman filter with the gain range 0.5-0.8. It is the output that is injected into the neuronal synapse. Below are all 16 possible binary permutations of APG output and hence βMN activation (but note that in the actual simulations both the amplitudes and unitary durations were also varied) with the corresponding names indicated along the x-axis. Gray backgrounds indicate permutations used to generate Figs 4 and 7.

**Figure 3.**
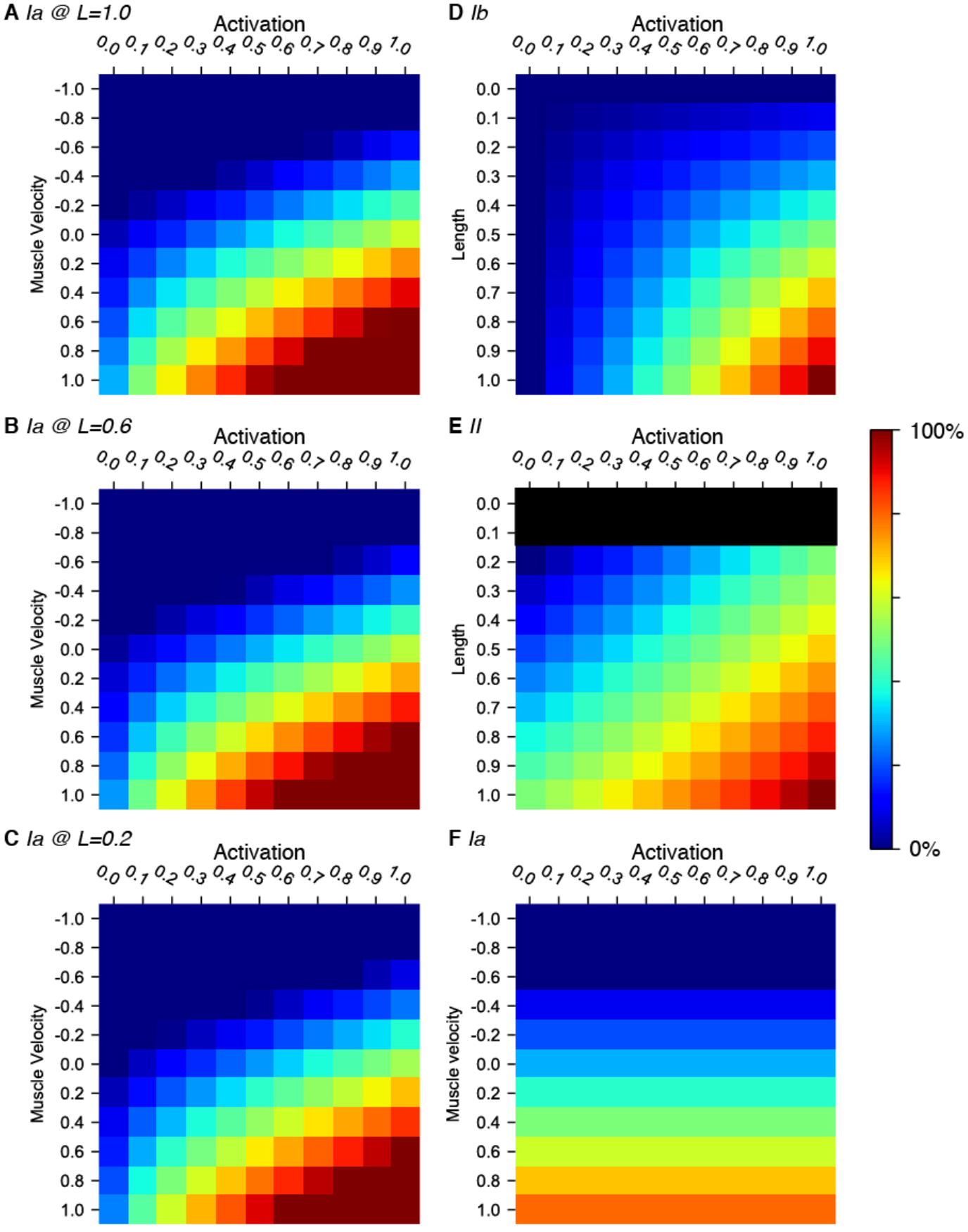
Output heatmaps illustrating the tuning of the sensors. A-C) Group Ia sensor output with muscle lengthening speed along the y-axis, and MN muscle activation along the x-axis. The II term of the Ia sensor activation (as defined in equation 6) varies with muscle activation, whereas muscle length is set at 1.0 in A, 0.6 in B and 0.2 in C. Muscle length 0.6 corresponds to the neutral position and 0.2 to the minimum length possible. D) Group Ib sensor output with muscle length along the y-axis and MN muscle activation along the x-axis. E) Group II sensor output again with muscle length along the y-axis and MN muscle activation along the x-axis. Note that the minimum muscle length possible is 0.2. F) Group Ia sensor output without fusimotor effect, thus only coding for muscle velocity and length. The muscle length is set to 0.6 in this heatmap.

**Figure 4.**
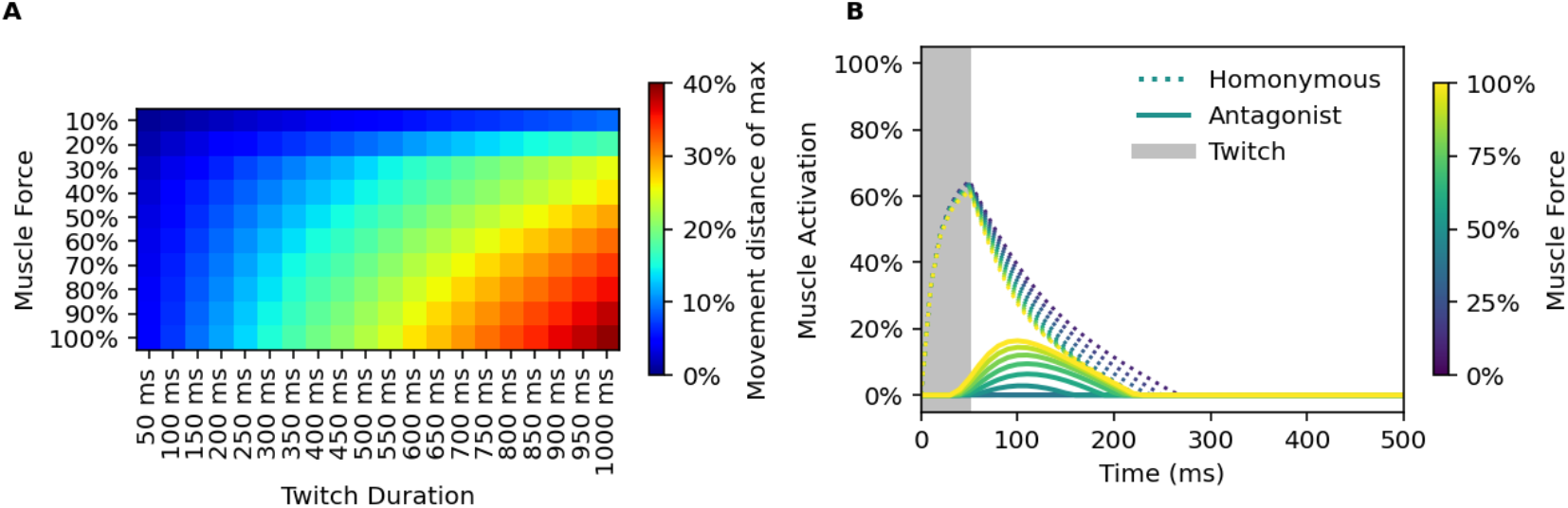
A) Relation between twitch duration (x-axis) and muscle force (y-axis) with respect to kinematic response. 100% kinematic response has been defined as movement from neutral position and to the full anatomical stop in one direction. B) Muscle activation responses due to a 50 ms 100% amplitude twitch into the homonymous MN with a connected NN with 100% synaptic weights for the homonymous Ia-MN synapses and 0% for the remaining, for iteratively increasing muscle force. A response from the antagonistic muscle is seen when the muscle force increases above 50% after training of the Ia-MN synapses.

**Figure 5.**
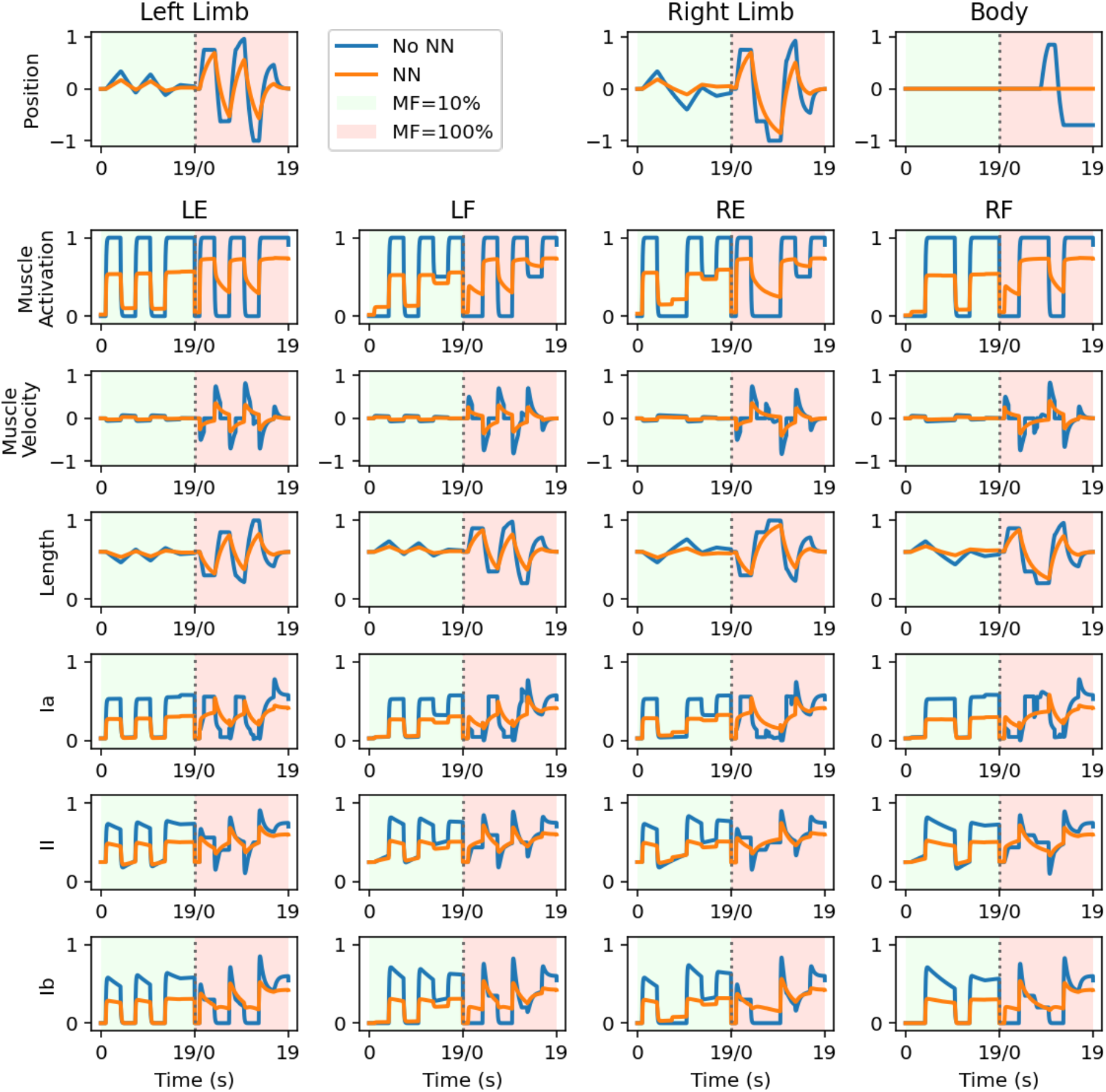
Plant dynamics with a duration of 19 seconds for a predefined set of muscle activations. During the first 19 seconds (green background) the muscle force of the plant is scaled down to 10% of max, and the second iteration (red background) the muscle force is set to 100% of max. First row plots relative position of respective limb and body with respect to their neutral position. Second row shows muscle activations for all muscles of the plant. Movements, as defined in Fig. 2C are Symmetrical extension, Symmetrical flexion, In phase leftwards, In phase rightwards, In phase leftwards with 50% contraction of the antagonists, and finally Full co-contraction. Blue lines indicate plant dynamics without any connected neural network. Orange lines indicate plant dynamics with a connected neural network. During 10% muscle force context the synaptic weights were at their initially randomized weights; during the 100% muscle force context the synaptic weights correspond to the mean end weights of Fig. 6A. The reason for the decline in all response amplitudes when a neural network has been connected is the leaky integration seen in equation 10.

**Figure 6.**
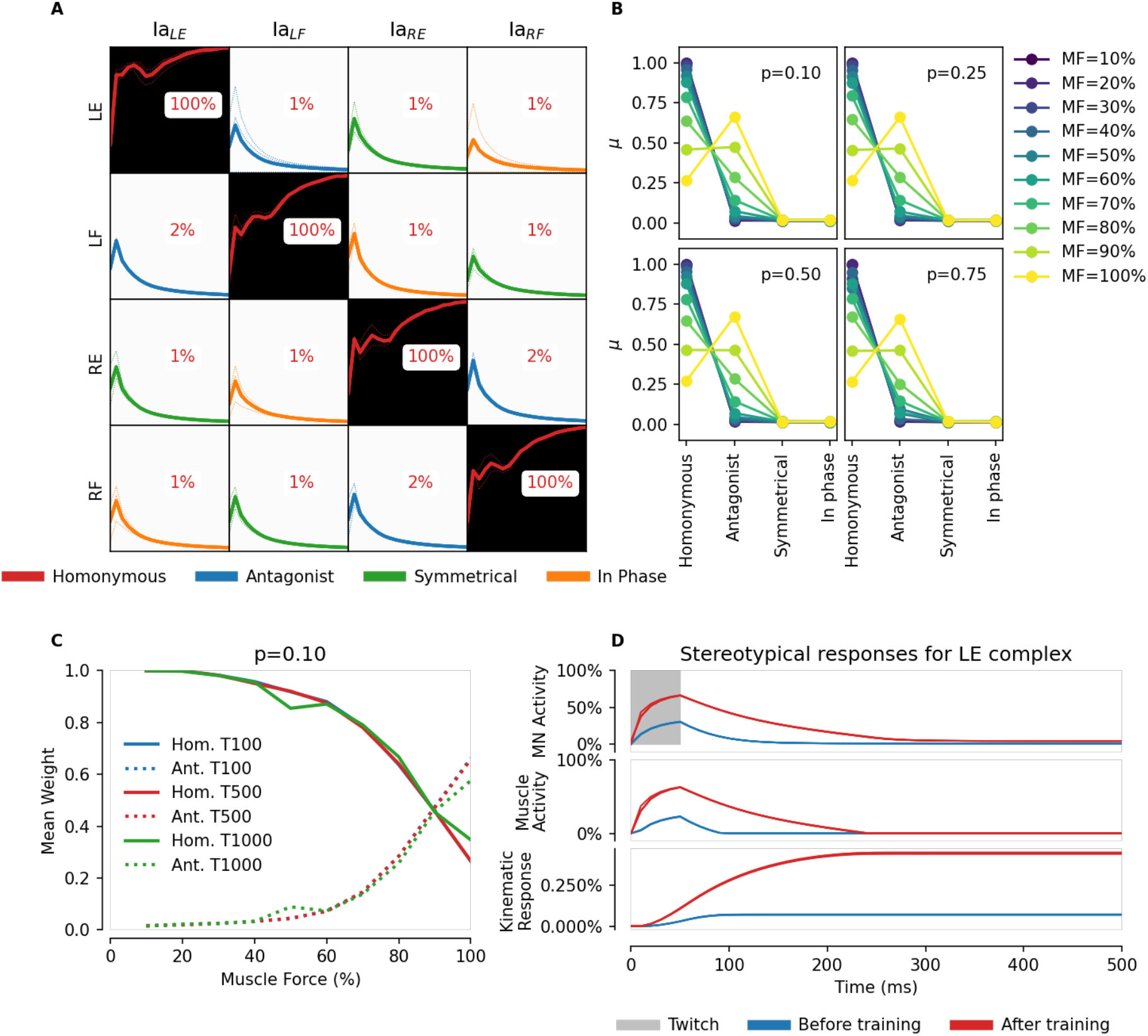
A) 4-by-4 synaptic weights matrix after training using the general APG. Showing individual simulation traces as dotted lines (N=3) and mean as solid line. Muscle force at 10% and Ia sensor with fusimotor innervation from its respective βMN. Each row illustrates one βMN, denoted by which muscle it innervates. Incoming Ia synapses are organized in columns, where the subscript indicates the muscle of origin of the sensor. Each cell in the matrix displays the mean of the temporal evolution of the synaptic weights for N=3 simulations with different randomized initial weights. The colors indicate the functional relationship between the βMN and the sensor according to the legend below. B) Summary of additional simulations with varying muscle force (10%-100% with 10% increments as shown by legend to the right). Each 4-by-4 synaptic weight matrix has been summarized by functional relationship between βMN and sensor as identified in panel A. Each category reports mean synaptic weight (N=3) as a data point. In each of the four panels the twitch probability used has been annotated. C) Mean end weight for homonymous (solid lines) and antagonists (dotted lines) for different muscle forces (x axis) and twitch durations (line colors). Twitch probability was p=0.1 for all simulations. Ia inversion is seen at 100% muscle force. D) Evolution of motoneuron activity and corresponding kinematic response. Traces procured from one of the simulations in panel A. Top plot shows left extensor motoneuronal response to a 50 ms square pulse twitch with additional feedback from Ia sensors. Middle plot shows respective resulting muscle activation of the left extensor. Bottom plot shows the kinematic response of the limb. The absolute amplitude of the response is small, but that is to be expected because the muscle force is set to 10% of max.

**Figure 7.**
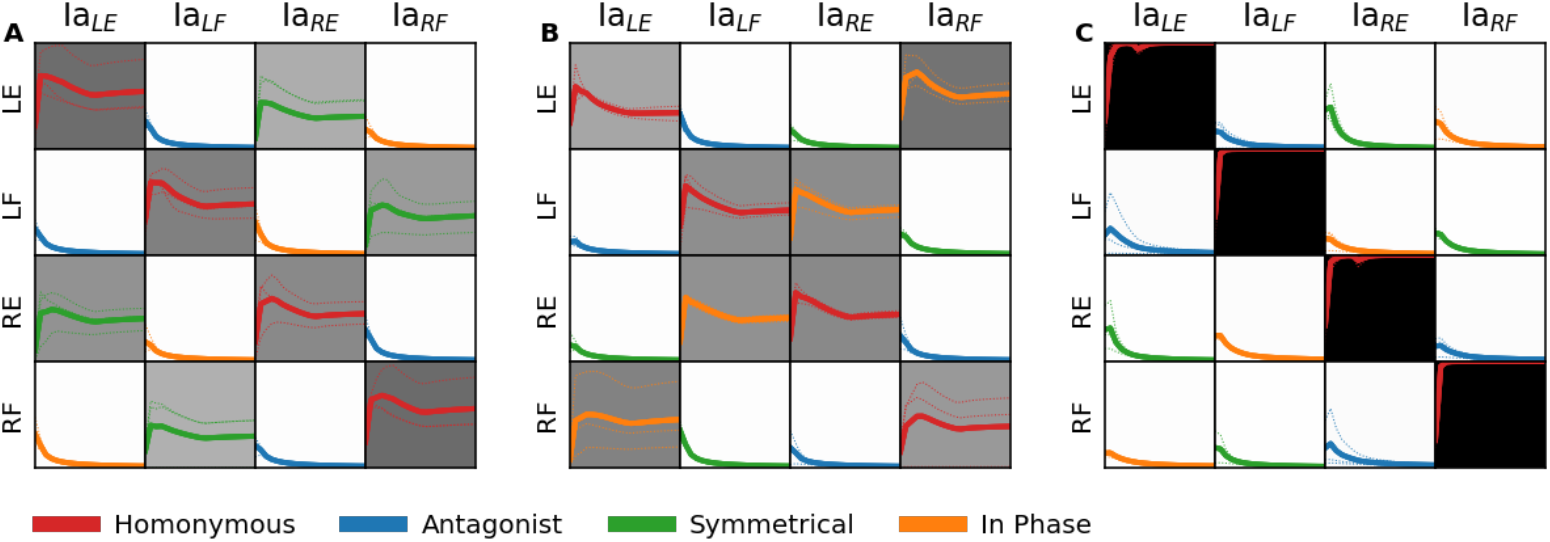
Synaptic weight matrices illustrating learning outcome induced after training with reduced APGs. A) The APG only generates symmetrical movements between the LE/RE and the LF/RF. B) The APG only generates in-phase movements. C) The APG never generates co-contraction. Each panel reports data from N=3 simulations, and individual traces as dotted lines. Solid lines indicate mean value. Twitch probability for all simulations was set to 0.1, muscle force was set to 10% of max.

In summary, the Oropod has limb and body dynamics dominated by velocity and muscle dynamics dominated by length (limb position), whereas real terrestrial organisms have limb dynamics dominated by acceleration and muscle dynamics dominated by length and velocity (Tsianos and Loeb, 2017).

### Somatosensation

Each Oropod muscle has proprioceptors that encode velocity, length and force generation (Fig. 2A), consisting of group Ia muscle spindle afferents, group II muscle spindle afferents and Golgi tendon organs (Ib), respectively. The response dynamics of these sensors during a predefined set of muscle activations (without any connected neural network) are illustrated in Fig. 5. Our model ignores other mechanoreceptors in muscles, joints and skin that contribute to proprioception. The Oropod Ib receptor is modeled as a simple, linear sensor of active force being generated by the muscle in which it resides (Fig. 3D) (Mileusnic and Loeb, 2006, 2009). The spindle receptors (Ia and II) are discussed in more detail below. The present study covers an early stage of fetal development in which only the Ia sensor has functional projections to the spinal circuitry (Eyre et al., 2000).

#### Beta Motoneurons for Fusimotor Drive

As noted in the Introduction, the natural tendency of extrafusal muscle fibers to shorten and thereby silence the spindle stretch receptors argues against the feasibility of Hebbian learning as a basis for the homonymous monosynaptic stretch reflex. However, amphibian and fish muscles are generally innervated by βMNs that simultaneously activate extrafusal and intrafusal muscle fibers (Eyzaguirre, 1957). Mammalian muscles also generally have a substantial but variable percentage of βMNs alongside their much more evolved and independent αMN and γMN subsystems that provide independent control of extrafusal and intrafusal muscles fibers, respectively (Manuel and Zytnicki, 2011). The fusimotor effects of βMNs are likely to be weak in adult mammals. βMNs probably fire at the same 10-50 pps as αMNs (Hoffer et al., 1987) whereas γMNs generally fire at 50-200 pps (reviewed by Ellaway, Taylor, Durbaba (2015)). During early fetal development, however, extrafusal muscle fibers are immature and weak (Gokhin et al., 2008), γMN activity may be absent or uncoordinated (Shneider et al., 2009), and the body tends to be confined *in utero* or *in ovo*. In this situation, the fusimotor effects of activating a βMN would likely produce a net increase in firing of the afferents in the spindle that it innervates. Thus, the βMNs could potentially support a learning-based organization of the Ia-MN connectivity. All or most αMNs and βMNs in a given motor nucleus are likely to be corecruited as a result of fetal tight junctions among them (Kiehn and Tresch, 2002).

#### Spindle Receptors and Fusimotor Function

It is important to consider the complexities of spindle receptor and fusimotor function that are largely preserved across vertebrate phylogeny (Matthews, 1974; Mileusnic et al., 2006). There are at least two distinct types of intrafusal muscle fibers, which have different mechanical effects on the sensory receptors that wrap around them. The group II receptor wraps around and detects simple length changes in fast-twitch intrafusal fibers (chain and bag2 types) whose fast-twitch contractions when activated can overcome shortening produced by extrafusal muscle, resulting in increased group II firing at a given muscle length. The Ia receptor is wrapped around all of the intrafusal fibers, but its location on the bag2 fiber is in the middle region where the bag of nuclei displaces myofilaments. Active contraction of the bag2 poles results in much larger biasing effects on the Ia than II receptor. The Ia receptor alone wraps around the compliant middle region of the bag1 fiber, a slow muscle type that becomes increasingly viscous at the poles with activation but does not shorten appreciably.

This results in a mechanical high-pass filter for externally applied length changes that accounts for activation-dependent velocity sensitivity of the Ia. These two types of Ia modulation tend to kick in at different levels of extrafusal recruitment. When mammals want to execute large, rapid movements, they recruit static βMNs that activate “fast” muscle fibers both extrafusally (fast-twitch = type II) and intrafusally (bag2 and chain). The intrafusal contractions result in “static” increases in activity of both spindle Ia and II afferents, essentially a positive bias that will keep these receptors from being completely unloaded and silenced during the large, rapid shortening of the whole muscle that is expected from the strong extrafusal muscle contraction. When mammals want to use muscles only to stabilize existing postures, they recruit only the “dynamic” subset of their βMNs that innervate slow (non-twitch) intrafusal muscle fibers (bag1) that enhance velocity sensitivity and slow and weak extrafusal fibers (actually slow-twitch or Type I in mammals) (Luna et al., 2015). Finally, adult mammals are equipped with two types of γMN (static and dynamic) that the CNS activates independently of the α and βMNs to provide differential control of length and velocity sensitivity of spindle afferents (Loeb, 1984; Mileusnic et al., 2006); γMNs differentiate postnatally (Shneider et al., 2009) and their effects neonatally are unknown.

We have modeled the Oropod muscle spindles as being all-beta innervated, like in amphibian systems whose anatomy but not physiology is well-described, while using the well-described and modeled physiology of the mammalian spindle to combine both static and dynamic fusimotor effects.

All somatosensory afferents generate excitatory post-synaptic effects centrally that are scaled and clipped, if necessary, to have a dynamic range that is 0 to +1. The ranges of possible lengths and velocities of the Oropod muscle are constrained by its limb mechanics, so the internal coordinates for each muscle can be normalized such that each muscle length (*L*) has the range 0 to +1 and each muscle velocity (*V*) has a range from -1 to +1 (Zajac, 1989). Muscle activations (*A*) and all sensory afferents range from 0 to +1.

Assuming the above normalizations, we define the sensitivity of the Oropod spindle receptors in the following sections.

#### The type II Muscle Spindle Afferent

The length-related sensor is the type II (secondary) muscle spindle afferent and its output is defined in equation 6 (and visualized in Fig. 3E). Its properties provide one term of the Ia sensitivity by virtue of their common receptor endings on the chain and bag2 intrafusal fibers. The *A* term provides the static fusimotor effect that scales linearly with extrafusal activation. Thus, the sensor would reach its maximal value of +1 only when the muscle is maximally stretched and maximally activated, e.g. during the reversal of a maximal waving motion.

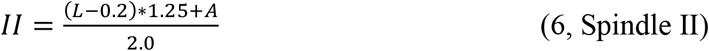

#### The type Ia Muscle Spindle Afferent

The velocity-related sensor is called the type Ia (primary) muscle spindle afferent and its output are defined in equation 7 (and visualized in Fig. 3A–C). Its muscle velocity sensitivity depends nonlinearly on muscle activation so that it increases rapidly for lightly recruited muscle and then saturates, simulating the early recruitment of the dynamic βMNs. Its length sensitivity is similar to the spindle II but with an added activation-dependent term reflecting the strong effect of wrapping around the middle region of the bag2 intrafusal muscle fiber.

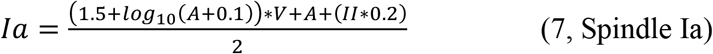

The maximal velocity of stretch of an Oropod muscle occurs when it is passive at its shortest length (A=0, II=0) and its maximally activated antagonist is at its longest length. This will generate output Ia=+0.25. In a maximally activated muscle (A=1), maximal velocity of stretch (V=+1) would saturate Ia at max (which is clipped by the range of 0-1). That condition can never arise without external interference because turning the muscle on necessarily decreases its rate of stretch by the antagonist. A maximally active muscle that is shortening at maximal muscle velocity (V=-1) would saturate at Ia=0 (which is physiological in mammals). A muscle that is active at A≈0.22 could shorten at maximal V≈-0.22, which would result in output Ia=0. Further increase in muscle activation or obstruction of shortening from antagonist or external load would result in a strong velocity signal (which is physiological in mammals).

For our control experiments, we also created a sensor devoid of all fusimotor effect, simulating a context with only αMNs (equation 8; visualized in Fig. 3F).

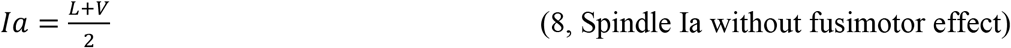

### Neuronal model design

To evaluate our hypothesis of self-organization of sensorimotor function based on musculoskeletal mechanics and behavioral experience, we needed an artificial neuronal model suited for repeated simulations of arbitrary neural network (NN) connectivity among a variety of afferent and interneuron types. To this end, we used a linear summation neuron model with dynamic leak (Rongala et al., 2021) together with learning rules similar to previous simulations of cuneate neurons (Rongala et al., 2018), i.e. Hebbian-inspired calcium co-variance learning rule.

#### Network Connectivity

Each of the four Oropod muscles is controlled by a single MN and has three proprioceptors that project to the NN (Fig. 2A). In this paper, however, we are mainly concerned with the Ia primary afferents that project directly to all four MNs (Fig. 2B).

In biology, muscles are not commonly controlled by a single MN alone. Rather, motor pools made up of hundreds of MNs control the activation of each muscle, thus enabling the muscle to be gradually activated by cumulative recruitment of motor units. In early development, the activities of these MNs are synchronized and proportionally recruited by means of tight junction coupling (Kiehn and Tresch, 2002). Based on this observation, and for purposes of model simplification, we here reduced the motor pool of each muscle into a single MN with a scalar output (*A*) with the functional range 0-1. However, due to the nature of the neuron model the hyperpolarized state of each neuron depends on the presence of inhibitory neurons. In this paper we do not have any inhibitory neurons and therefore the neurons would only be in their dynamic active range. To compensate for this we added a bias to the output activity from each MN such that:

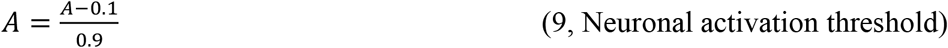

#### Neuron Model

Physiologically, our neuron model is electrotonically compact in the sense that the synaptic voltage signals are added linearly to the somatic voltage. In our model we have two categories of ‘biochemical’ compartments, a main, somatodendritic compartment and synaptic compartments, where the compartmentalization serves the sole purpose of controlling the learning. This approach has been used to model learning in cuneate neurons (Rongala et al., 2018). Cuneate neurons, however, have relatively complex calcium dynamics in their main compartment that the synaptic inputs can trigger. Here, we simplified that model to instead make the calcium concentration (as a dimensionless quantity) of the main compartment equivalent to its voltage (which in turn was defined by a linear summation of the synaptic inputs).

#### Neuronal Compartment Model

The output of most CNS neurons is the action potential. However, in our previously published (Rongala et al., 2021) neuron model spiking is omitted, and each signal between neurons is instead a time-continuous voltage signal (*A*), which can be thought of as representing the combined activation of a population of asynchronously firing neurons of a given type. This simplification is valid because the spike output of the neuron can be considered a somewhat noisy probability density function of its membrane potential (Spanne et al., 2014a). The discreteness associated with the timing of single spikes in individual sensory neurons can be assumed to be mostly averaged out across the population of asynchronously firing neurons of the same type (Spanne et al., 2014b).

Signal integration in neurons depends on the establishment and the modulation of the membrane potential. The resting membrane potential depends on the background leak conductance, which is dominated by potassium. Modulations of this membrane potential are generated by the activation of chemically or electrically gated conductances for specific ions (i.e. in excitatory synapses the chemically gated conductances are typically dominated by sodium). Action potentials are generated with a frequency that is more or less linearly related to the degree of depolarization (Spanne et al., 2014a).

Background excitatory or inhibitory synaptic activity itself reduces the response to any particular synaptic input because it is accompanied by opening conductance channels in the postsynaptic membrane (Bannister et al., 2002; Sahara and Takahashi, 2001; Silver et al., 1996; Traynelis et al., 1993) and thus results in shunting. This is particularly important in a developmental model of the NN in which the number and strength of synaptic inputs onto any given neuron is expected to undergo large changes (see also Tsianos et al., (2014)).

Rather than simulating the conductances explicitly, these effects are instead emulated using equation 10, where A_i_ is the output from the *i*th neuron, which depend on the input from *j*th neuron (A_j_), modulated by the synaptic connection weight (w_ij_) and a sign matrix where excitatory synapses are positive and inhibitory are negative (S_ij_). The shunting is represented by the sum of synaptic activation plus a resting leak constant (k_i_) in the denominator which is the rolling sum of synaptic activation (x) as obtained by equation 11 (K is the gain factor). The summed synaptic activity in both the numerator and in the denominator are low-passed filtered using a Kalman filter as described in equation 11.

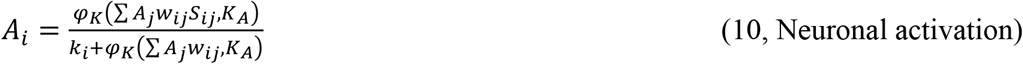

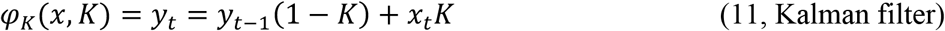

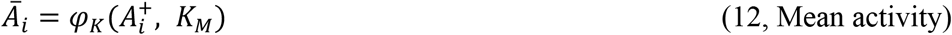

This neuronal activation behavior has the important feature of protecting each neuron from saturation during development, as each active synapse adds to the activation with a potentially increasing weight, but then also increasingly adds to the leak.

Equation 10 provide a complete description of how information flows through our neuronal model. The output from any network configuration composed of these neurons is fundamentally dependent upon the particular synaptic weights.

#### Synaptic Weights

The synaptic weights in our model are scalars ranging from −1 to +1. There is no information on fetal synaptic weights, and thus we made the assumption that synapses have randomized initial weights at the low end of the spectrum. In our model the initial synaptic weights are random samples from a normal distribution (mean μ=0.2, standard deviation σ=0.16, synaptic weights below 0.001 are reseeded thus rendering all initial weights positive). The model can in theory reduce the synaptic weights into the negative range, but only through learning.

#### The Dynamics of the Learning Mechanism

During the course of a simulation, the synaptic weights are continuously adjusted according to the correlation between the activity of the individual synapses and the activity level of the neuron as a whole. This learning mechanism is in essence the calcium co-variance rule (Dean et al., 2010; Sejnowski, 1977) implemented such that both Hebbian LTP and LTD of synaptic plasticity can occur. The core learning rule is an adaptation of Oja’s rule (Oja, 1982) and is described in equation 13. The main difference between this equation and Ojas rule is that instead of a binary parameter that allows for update or not, a scalar learning signal parameter has replaced it (denoted *l*, equation 14, subscript *ij* indicates the jth synapse of the ith neuron). The dynamic that governs the learning signal has several important aspects, but essentially the degree of correlation between the activity of the synaptic compartment and the neuronal compartment (equation 14) is what will define the fate of the synaptic weight. This learning signal is a scalar parameter in the range of −1 to 1, and thus allows for scalable potentiation and depression of the synapse in question. Depending on the sign of the learning signal, and thus the direction of update, the compensation factor (c) is changed according to equation 15. Finally, a learning rate parameter (ηi for the ith neuron, equation 16) scales learning.

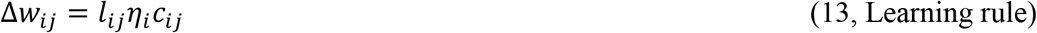

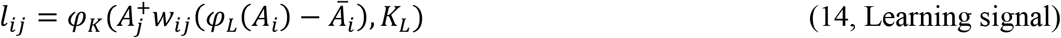

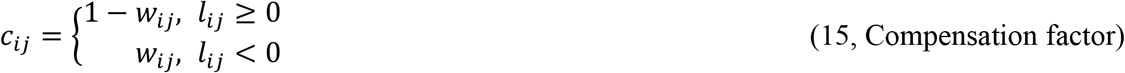

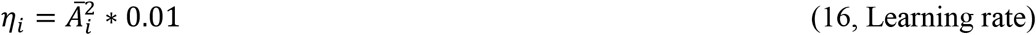

#### Learning Signal

The mean potential (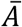 in equations 14 and 16) is meant to encompass a number of more complex features of the internal machinery of the neuron. First and foremost it is meant to model the current balance of phosphatases and kinases in the neuron (Jörntell and Hansel, 2006), which is assumed to be controlled by the total calcium level, which in the present model is simply the output activity generated by the neuron over time. A higher concentration of kinases, for example, can make the probability for LTP higher, whereas a higher concentration of phosphatases would instead increase the probability for LTD. A high probability of LTP will over time result in a higher output activity of the neuron, in which case our model neuron would respond by shifting the balance towards phosphatases to prevent overexcitation. An ongoing adaptation of this polarity threshold hence helps preventing saturated learning in the neuron (Rongala et al., 2018) and in the current system with the synaptic input from the sensors being spontaneously active it has the net effect of implementing homeostatic synaptic plasticity, or synaptic scaling (Turrigiano, 2017).

### Autonomous Activity Pattern Generator (APG)

In any organism or model thereof, a central issue is how activity initially arises and how it is subsequently molded into increasingly useful behavior. In the mature spinal cord the notion of a central pattern generator (CPG) is ubiquitous (for a review see Guertin, (2009)). In many envisaged instantiations, the CPG has the power to produce functionally synergistic contractions of muscles, and thus produce movements that are functional, hence evolutionarily sound. However, it is still not clear how or when the formation of the CPG would reach such a stage of maturation. Here we chose the more assumption-free approach of letting the MNs generate their own excitation internally. This approach seems to be supported by evidence for pacemaker potentials in MNs arising from a high level of voltage-gated calcium channel expression and resultant calcium positive feedback responses in early developmental stages (Corner, 1977; Tabak et al., 2015). These and other intrinsic conductances support the generation of pacemaker-like potentials in individual MNs. Given the high prevalence of gap junctions between the MNs of a given motor pool early in development (Kiehn and Tresch, 2002), such pacemaker activity output could recruit large numbers of MNs synchronously and proportionally, which would account for visible limb twitches in fetal animals. In our model system, such autonomous activity pattern generation is assumed to synchronize across the MNs of each muscle. The Activity Pattern Generator (APG), hence consisting of internally generated pacemaker potentials initiated at random in the single MN pools, was designed to activate the MNs of separate muscles independently in random patterns.

#### APG patterns generated in the simulations

Each APG pacemaker potential has four parameters: amplitude, duration, rise time and probability of activation. Each parameter was set in order to avoid biasing the model towards any particular solution in the absence of systematic experimental data about these parameters.

The amplitude of activation was assigned a random value between 0 and 1 from a uniform distribution. Since we employed the simplification of modelling the entire MN pool as a single neuron we had to let that particular neuron be able to express the full range of activation amplitudes.

The duration of each APG potential was used to mimic the observed short muscle twitches in the embryological stage (Corner, 1977). For the primary set of simulations, the duration of each APG potential was randomly assigned a value between 50 ms to 100 ms from a uniform distribution. To assess the importance of these twitches as opposed to sustained phasic movements observed during later fetal stages, we made additional simulations with longer duration APG potentials. The kinematic effect on the homonymous muscle of varying twitch amplitudes and durations can be seen in Fig. 4A. After maturation of muscle force and learning in the Ia-MN synapses, even the shortest duration twitches (50 ms) produce limb movements that are large enough to produce a reflexive response from the antagonist muscle (see Fig. 4B and Results).

To soften the *a priori* assumptions, we also applied a one-dimensional Kalman filter to each of the generated APG potentials with a gain randomly assigned a value between 0.5 to 0.8 from a uniform distribution. This had the effect of smoothing the otherwise square-pulse inputs (Fig. 2C).

Finally, the probability of activation is equal to the notion of frequency of muscle twitches. This feature of the APG potentials will have two main impacts to be noted. First, since the generation of APG potentials for each MN are independent, an increase in probability of activation would increase the probability of two, or more, MNs to be active at the same time. This is important because the existence of four MNs results in 16 different permutations of activation (Fig. 2C), each of which has the potential of exhibiting different dynamics of the plant in different contexts. For example, strong extensor muscle activity starting from a neutral limb position results in very different dynamics as opposed to if it starts from an extreme end-position. In a more complex plant this would include co-activation of synergists. Because each APG potential is independent, however, an increased overall probability of activation increases the probability for co-activation of non-functional MNs. If a non-related MN was consistently co-activated with a particular MN, then a Hebbian learning rule would strengthen the synaptic projections between them. Furthermore, this potential problem of spurious connectivity will be less prevalent in a plant with more muscles since the probability of repeated co-activation would decrease. Previous work has touched briefly on this topic (probability of activation set to 11% in Peterson (2003)), but we here explore it further.

The purpose of our choice of setup of the general APG was as follows: Random activation of MN pacemaker potentials between muscles should eventually, or during the course of development, result in exploration of every combination of muscle activation in every posture. These will include instances where limb movement is impeded or obstructed by coactivation of antagonist muscles, by contact between limbs or by contacts between a limb and a wall.

We also created variants of APGs that were limited to subsets of the general APG, which might be generated by a central pattern generator (CPG) composed of genetically hardwired interneurons (such as suggested by Lundberg (1981) and Grillner (1985)). These variants include i) co-contractive activations are prohibited, ii) only symmetrical movements are allowed, and iii) only in-phase movements are allowed.

#### APG implementation

In the current model system, the APG was implemented as an excitatory synapse on the MN. The weight of that particular synapse was set to the max allowed weight (1.0) and was not allowed to learn. The amplitude of the excitatory drive from the APG summed with and had its effective strength modulated by all other synaptic inputs to the MN.

### Simulation

The nervous systems of real organisms are closed loops where the output affects the plant in some way, and these effects together with changes in the physical world create the sensory feedback that is projected back into the nervous system. The continuous sensory feedback is then mixed with internal signals and produces an updated output from the nervous system. We emulated this loop with our artificial plant (see initial part of Methods), our artificial nervous system (see Neuronal Model Design for details), and an artificial physical world as follows:

1. Whenever an assigned state of the APG had elapsed, the APG was given a new state with a random activation amplitude, duration and rise time as described above.
2. The time evolving activity of the Ia sensors were combined with the current APG state in each MN as described above. Hence, for each MN there was a time-evolving output level on which the learning process elaborated the synaptic weights.
3. The contractile force generated by each muscle in the plant was updated according to its current length and the output from each corresponding MN, thus updating the state of the plant.
4. The state of the Oropod plant and its interaction with the surrounding physical world was continuously updated by the physics engine (pymunk, http://www.pymunk.org/) using an integration time step of 10 ms.

We saved the state of the NN continuously throughout our simulations. This allowed us to load any saved state and run analysis on that particular state. Each simulation ran for 20000 seconds, which was a duration that allowed synapses that were initially set to a very low value to adapt to a stable higher value with a wide margin.

We could load a NN state into the loop described above and let it perform a specified set of movements. This was achieved by replacing the APG with a profiling module that produced a known set of activation states (defining which MNs to activate also with respect to amplitudes and durations; Fig. 5).

## Results

We have designed a model creature with a simple set of musculoskeletal mechanics and biologically derived sensor tunings (Fig. 1 & 3). The model creature was controlled by a neuronal network with biologically derived synaptic learning rules and a simple, randomized activity generator in order to test the feasibility that some main features of adult spinal cord circuitry connectivity could be explained by learning-driven self-organization (Fig. 2).

### Random MN activations promote spinal connection patterns

The synaptic weight learning of our first set of simulations with the general APG (N=3, each made with different initial synaptic weights between the Ia sensors and the βMNs, 10% muscle force and twitch probability of 0.1) quickly converged such that Ia-afferent feedback synapses for the homonymous muscle were potentiated whereas those from the non-homonymous muscles were depressed (Fig. 6A, the grand mean synaptic weight of the diagonal was 1.0, std=7.5e-6), i.e. a similar synaptic weight matrix as observed for Ia-MN synapses in adult mammals (Eccles et al., 1957). To evaluate the impact of such severely reduced muscle force we ran additional simulations with incremented muscle forces, increasing by 10% up to the maximum of 100% (N=3 for each setting). Not surprisingly the specificity of the homonymous Ia synaptic strength started to decline as the extrafusal muscles gained in strength. The mean homonymous Ia-MN synapse weight remained at a high weight until a level of around 60% muscle force where a sharper decline started. At 90% muscle force the antagonistic Ia-MN synapse weights were equal to the homonymous; a reversed connection scheme, where the antagonistic Ia-MN synaptic weight was higher than that of the homonymous, arose at 100% muscle force. The increase in weight naturally produced a larger kinematic response (Fig. 4) and the kinematic response reduced the tension of the intrafusal muscle spindle, reducing the feedback on which the Hebbian learning depends (Fig. 6B).

Both the frequency and duration of the twitches will modulate the kinematic response. We re-ran the above simulations with the twitch probability set to 0.25, 0.50 and 0.75 (Fig. 6B). However, the results were near identical to the set of simulations with a twitch probability of 0.1, so twitch probability did not affect outcome. Next was the duration of the twitches, where a longer twitch would in theory give the muscle longer time to unload the muscle spindle. To explore this, we re-ran the above simulations once again, but with twitch durations of 450-500 ms and 950-1000 ms. The twitch probability was in all simulations set to 0.1 (Fig. 6C). In summary the twitch duration did not alter the initial outcome seen in Fig. 6A, B. This indicates that the precise nature of the muscle twitches did not play a crucial role in the outcome of the synaptic learning, whereas the relative strengths between intra-and extrafusal fibers did (Fig. 6B).

We also evaluated the behavior of the system before and after learning. In the untrained network, a 50ms square pulse at 100% twitch strength was injected into the left extensor motoneuron. The injection produced neuronal response as well as movement (Fig. 6D, blue lines). After training, however, the same pulsatile injection produced a much more elongated neuronal response with a correspondingly elongated kinematic response of the limb (Fig. 6D, red lines). Hence, training increased the capacity of the network to sustain movements long after the termination of the movement command. When tested with the profiling APG pattern before and after training, this effect resulted in stronger muscle impulses and more complex limb dynamics (Fig. 5), including the beginnings of oscillatory limb movements due to stretch reflexes (antagonistic response dynamics can be seen in Fig. 4B).

### Reduced APG results in less distinct connection patterns

While our APG has a conceptual relation to the CPG theory, CPG theory promotes the idea that there exists pre-wired neuronal circuitry that produces behaviorally relevant cyclic alternating activations of muscles that could support locomotion. Hence, by this definition a CPG would generate only a subset of the theoretically possible MN activation combinations. Therefore, we also tested reduced variants of the APG where not all theoretically possible MN activation combinations occurred, with muscle force set to 10% as above. Training with the APG variant that produced only in-phase movements (see Fig. 2C and Fig. 7A) resulted in a moderate homonymous Ia-MN connectivity (N=3, mean=0.41, std=±0.044) plus an equal connectivity with the in-phase synergist in the contralateral limb (mean=0.48, std=±0.045). Conversely, the APG variant that produced only symmetrical movements (Fig. 2C and Fig. 7B) again resulted in a moderate homonymous Ia-MN connectivity (N=3, mean=0.53, std=±0.044) and was in this case also accompanied by a moderate Ia-MN connectivity (0.36, std=±0.05) between the extensors of each limb and between the flexors of each limb.

A less reduced variant of the APG, in which only muscle co-contraction was forbidden, produced a synaptic weight pattern (Fig. 7C) that was similar to that of the non-reduced, general APG (Fig. 6A). The homonymous projections (diagonals) were clearly potentiated (N=3, mean=1.0, std=±6.4e-14) and the remaining synaptic connections pruned.

### The fusimotor effect on the Ia sensors is necessary for learning

As a control, we eliminated all fusimotor effects of the MNs, effectively simulating an organism with only αMNs. With this adaptation of the Ia sensor, we ran a set of simulations (N=3) with the muscle force again set to 10% and twitch probability 0.1. In this case, the synapse-specific potentiation of the Ia-MN projections seen in previous simulations disappeared (Fig. 8A). However, the only way to get a reasonable response from any Ia sensor in this case is through overt limb motion. This would necessitate a higher muscle force, thus we ran an additional set of simulations (N=3) with the muscle force set to 100% (Fig. 8B). In this case the homonymous synaptic weights were depressed rather than potentiated and there was some potentiation of the Ia synapses on the antagonists. Interestingly, however, there was still extensive learning in the NN, but in this case all the synaptic weights were driven towards zero. We confirmed that the plant kinematics during APG input was largely preserved (Fig. 8C). In fact, all Ia sensors had substantial activity during the training but these activations were not well correlated with the activation of the muscles.

**Figure 8:**
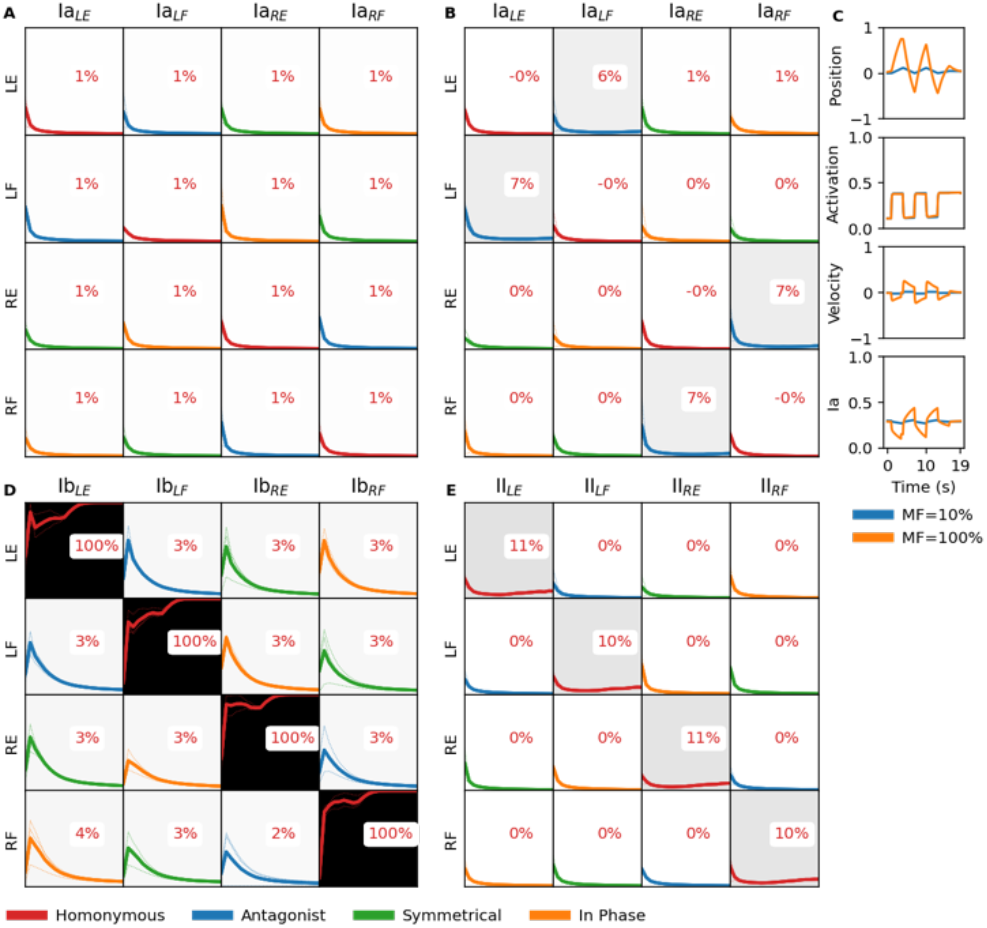
Learning outcomes after training using supplementary contexts. A) Ia sensor is left intact but the fusimotor of the motoneurons has been turned off, thereby simulating αMNs instead. Muscle force was set to 10% of max, twitch duration 50-100 ms with a probability of 0.1. B) Equal to context in panel A but with muscle force set to 100% of max. C) Example traces for limb position, muscle activation, muscle velocity and Ia-activity for a set of predefined movements (equal to first column in Fig. 5) but using αMN as in panels A and B. D) Ia sensor feedback has been changed to Ib sensor feedback. E) Ia sensor feedback has been changed to II sensor feedback.

### Non-biological sensor input and Hebbian learning

For curiosity we also tried exchanging the Ia sensors with the Ib and II sensors, even though neither of these proprioceptors are known to make monosynaptic connections to motoneurons. We ran N=3 simulations for each variant while keeping the remaining parameters the same (muscle force at 10%, twitch duration 50-100 ms, twitch probability at 0.1). The simulations using Ib quickly converged to a strong homonymous synaptic strength, while decorrelating the remaining inputs. This is not surprising since any tension in the extrafusal fibers will compress the Golgi tendon organs and thus generate sensory feedback which can drive the Hebbian learning. This homonymous relationship between motoneuronal activity, muscle contraction and direct homonymous sensory feedback will for the Ib sensors in fact accelerate as the extrafusal fibers increase in strength, as opposed to the Ia sensor activity that will eventually invert (Fig. 6B, C) due to increasing negative muscle velocities.

In contrast, the simulations using group II as the sensor input did not end up with a strong homonymous connectivity but stayed low throughout the simulation. This showcases an interesting aspect of the neuronal learning model where it seems to be able to find the graded connection in a moving target. In this case, the stronger the output from the motoneuron the less the group II input will be, since a larger kinematic response equals a larger decrease in muscle length, which is what the group II monitors. However, there is still a correlation between the motoneuron activity and the group II input, since the latter is dependent on the output from the motoneuron (see equation 6).

## Discussion

### Nature vs. Nurture

The present study provides a basis for functional self-organization of the Ia-MN connectivity in the spinal cord based on sensorimotor experience resulting from random activations of these MNs during early development (i.e. nurture). The selective projection of Ia afferents onto the MNs of their homonymous parent muscle did not require any *a priori* knowledge of which MNs innervated which muscles. Instead, it emerged from the random activation of the musculoskeletal anatomy, but only when that included co-innervation of both intrafusal and extrafusal muscle fibers, as provided by βMNs (as opposed to independent α and γMNs). These results are in line with studies into myotonic specificity where recovery of functional behavior occurred after cross-connection of peripheral nerve fibers (Arora and Sperry, 1957). Indeed, βMNs are known to be widely present in mammals (Illert, 1996), whereas mature γMNs are present only in adult mammals and only gradually develop from perinatal stages and onwards (Ashrafi et al., 2012; Shneider et al., 2009). Hence, the effects observed here are likely to be dependent on βMNs. However, similar effects COULD be achieved if α and γMN activations were tightly coupled by gap junctions at this early stage of fetal development. It is worth noting that α and βMNs are likely to be so-coupled, which would result in homonymous spindle afferents onto both α and βMNs in a given motor nucleus.

The developmental role for βMNs proposed here begs the question of how the somewhat weaker but still common heteronymous Ia-MN synapses form between synergistic muscles. The Oropod is not suited to explore this due the absence of synergistic groups of muscles. Nevertheless, the results presented here suggest that a strong muscle twitch into a muscle would unload both the homonymous and synergist spindles; without the driving βMN input the synergists would fail to potentiate the appropriate synapses. However, along the same line of reasoning, a similar twitch into an antagonist muscle would introduce a positive muscle stretch in all of the synergists, thereby activating their Ia afferents. If their homonymous synapses resulted in reflexive activation of their MNs (as demonstrated in Fig. 4B), the simultaneous afferent and efferent activity among the synergists could potentiate the heteronymous Ia-MN synapses. Note that this could happen only at a developmental stage when the growing strength of the extrafusal muscle fibers starts to produce overt limb motion (see Fig. 6C). Such motion would enable the development of the fairly widespread interconnectivity among partial synergists (Eccles et al., 1957; Fritz et al., 1989) to reflect their functional, as opposed to anatomical, relationships in the limb.

Our results indicate a dependence upon the interplay between the relative strength of extrafusal and intrafusal muscle fibers, which is likely to shift during development. For example, at birth, or hatching, terrestrial animals must suddenly deal with gravity. Given that skeletal muscles become stronger with exercise, we made the assumption that in early development intrafusal fibers are ‘stronger’ than extrafusal fibers (Gokhin et al., 2008), i.e. whenever there is βMN activation, the Ia sensors hence more active, despite the unloading effects of extrafusal force generation. This is a necessary phenomenon for the learning we describe here to work, which will require future experimental testing. Another prediction from our model is that this phenomenon is probably dramatically reduced over time after birth, during which time extrafusal fiber strength has indeed been observed to increase tenfold or more (Gokhin et al., 2008). At that point in development, however, the Ia-MN synapse may be less plastic and/or the γMN system may start to provide the adult patterns of alpha-gamma coactivation that tend to accompany behaviors in which the muscles are expected to shorten (Loeb, 1984).

Another aspect that might have a high reliance upon critical periods of allowed plasticity in forming synapses is the connectivity of Ib fibers. When substituting the Ia stretch sensor with the Ib force sensor in our model system, the resulting connectivity matrix was as specific (Fig. 8D) as for the Ia (Fig. 6A). Nevertheless, monosynaptic Ib connections to MNs are not found in adult animals. Notably, muscle spindles with equatorial Ia afferent terminals are found as early as E19-20 in the rat (Zelená, 1957), while the entanglement of Ib nerve endings in the collagen bundles of Golgi tendon organs (GTO) is not observed until the first post-natal week (Zelená and Soukup, 1977) and they continue to grow until week 3. The difference in adult connectivity could thus result from a critical period in the development of *any* monosynaptic afferent input to MNs instead of a genetic difference in the afferents themselves. In this context, it is worth noting that both of these Group I afferents appear to induce the formation of their respective, specialized mechanoreceptive structures in fetal muscles but they can cross-innervate the wrong mechanoreceptors following peripheral nerve repairs in adults (Mendell et al., 1987).

### Neuronal model

The neuronal model used in this article (Rongala et al., 2021) provides a simplified neuronal calcium dynamics model with a continuous output signal rather than spikes. Oja’s learning rule (Oja, 1982) that we adapted for our model generates synaptic potentiation or depression according to the relative timing of calcium transients in the pre-and post-synaptic compartments of the synapse. We assumed that the post-synaptic calcium concentrations reflected simply the level of activity of the post-synaptic cell, but this is not always the case. For example, the inherent calcium dynamics of cuneate nucleus neurons causes the post-synaptic calcium to reflect a time derivative of the underlying membrane potential (Rongala et al., 2018). If such a situation obtained in MNs, the learned synaptic modifications would depend on more dynamic aspects of MN activation; in the limit this would resemble a spiking neural model. Unfortunately, little seems to be known about dynamic calcium gradients in early embryonic spinal MNs, so we elected to use the simplest assumption.

Both the synaptic weight matrices resulting from the learning process and the end synaptic weights of the diagonal in these matrices were remarkably stable over a large range of initial weights (N=3; see Fig. 6A). This stability was also evident over a large range of muscle forces, twitch probabilities and twitch durations (Fig 6), and also when considering non-biological sensor input to the motoneurons (Fig. 8D, E). Considering the robustness of our neuronal learning model, the end results may be robust across a further range of implementations.

The low dimensional biomechanics and network we studied here can be seen as quite trivial in relation to the biological counterparts. Here we focused on illustrating the principle that self-organizing spinal cord circuitry could be made to work. Future studies are needed to show if the system could work similarly with a higher number of degrees of freedom in the plant, with partial synergists such as multiarticular muscles and/or with a larger network and with other sensory modalities and the interneurons that convey them to MNs.

### APG versus CPG

When we trained our Oropod model to subsets of the theoretically possible combinations of muscle activation, i.e. restrictions that one would expect if the activity of the controller was driven by a CPG rather than a random muscle activity generator, the resulting connection matrices altered with respect to some critical features. The most restrictive of the subsets included only symmetrical or in phase movements; in both cases these resulted in heteronymous synapses from the Ia sensors of the synergistically activated muscles.

The least restrictive CPG subset prohibited cocontractions and resulted in a Ia-MN synaptic weight matrix similar to those observed in adult mammals (Eccles et al., 1957; Hongo et al., 1984). A completely random APG activity in the fetal animal could gradually give way to CPG patterns that generally avoid any cocontraction. Furthermore, the strength of the unopposed extrafusal muscle increases relative to the impedance of the limbs, resulting in even greater shortening velocity in the agonist muscles. If the synaptic plasticity of the Ia-MN projections remained unchanged during such development, the specificity of the homonymous loops might decline. Such an outcome could be avoided by introducing a limited critical period in this synaptic plasticity and/or developing γMN activity whose intrafusal effects counter these kinematic effects. The timing of each of these hypothetical changes during fetal and perinatal development is likely to be critical for successful ontogeny, so might be expected to be embedded in genetically specified general rules for development. Failure to follow any single step in this choreographed ontogeny would likely give rise to a wide range of secondary changes in the circuitry that would be observed as pathological phenotypes in the adult.

### Comparative Development

If fetal and perinatal development depends on timed sequences of events, how much recovery will be possible in the event of injury to either musculoskeletal or neural components at various times of life? Extraordinary adaptation to supernumerary muscles and limbs has been observed in amphibia but appears to depend on age. Such adaptation occurs quickly in larvae but slower in older animals (Weiss, 1936). Even among species with similarly freely swimming larval stages, there are substantial differences in the consequences of muscle paralysis on subsequent behavioral deficits (Haverkamp and Oppenheim, 1986).

A counterpoint can be found in the fact that an embryo confined *in ovo* has a quite restricted space in which to move (Hamburger and Hamilton, 1951), resulting in a restricted posture of the embryo during development (Bradley et al., 2014). Such restrictions might have some influence on the outcome of development (Bradley et al., 2014), but the end result is still very much functional (Sun et al., 2018). Such restrictions do not negate that the outcome requires extensive learning, rather that the sensitivity to posture and movement may be somewhat different in these species. Thus, even though our APG in theory exposes our system to every combination of activations in every posture, which might be a richer experience than for many biological counterparts, the increased richness due to space and overt movement does not seem to be the main source of our results. Rather we see correct homonymous connectivity arising due to very low extrafusal muscle force (low relative unloading of muscle spindles) while not being disrupted by increased twitch duration (increased movement, Fig. 4).

In adult mammals, nerve crossings and muscle transpositions result in persistence of the original neural circuitry and function (Forssberg and Svartengren, 1983; Gordon et al., 1986; Sperry, 1941), whereas muscle transpositions in neonatal cats result in changes in at least some muscle function and cutaneous reflexes (Loeb, 1999). Congenital duplication of the forearm and hand in man has been described with relatively intact sensorimotor function (Weiss, 1935). Absence of sensory feedback during development in humans results in dysfunctional muscle use (Suster and Bate, 2002). All of these suggest that there are critical periods of development during which the spinal circuitry is sensitive to patterns of sensory input.

The particulars of the nature versus nurture debate seem to be obscured by the need to account for highly functional connectivity seen in the adult while making vague assumptions about the potential consequences of the details of the learning rules that might support a nurture mechanism. An illustrative example is the formation of specific monosynaptic sensory-motor connections in the absence of neural feedback (Mendelson and Frank, 1991), while at the same time exhibiting a twofold difference in amplitude and reduced selectivity when compared to a control. Some limited monosynaptic specificity might arise from anatomical gradients of trophic cues in the target mesenchyme (Wenner and Frank, 1995), while the more precise connectivity is later tuned by learning rules (Kudo and Yamada, 1987; Saito, 1979; Seebach and Ziskind-Conhaim, 1994). Hebb himself appears to have skirted the complexities of embryological development and their potential dependence on his eponymous learning rule (Oppenheim and Haverkamp, 1986). The persistence of at least some appropriate monosynaptic synapses in some species following the withdrawal of functional cues may simply reflect a Hebbian rule that is biased toward synaptic preservation rather than elimination.

What we have demonstrated here is that muscle-specific Ia connectivity *can* develop with appropriately configured type-specific rules, namely the presence of βMNs linked by tight junctions and prone to spontaneous twitches and a critical period of synaptic plasticity while the extrafusal muscles gain strength. In other work underway, we are exploring whether this general principle can be extended to the interneuronal circuits of the spinal cord.

Reliance on nurture rather than nature during ontogeny, wherein the physical properties of the biomechanical plant can be imprinted on the NN without any *a priori* knowledge, is potentially important to understand the successful evolution of new species. Such evolution is driven largely by mutations that initially affect musculoskeletal mechanics. An individual with a potentially useful musculoskeletal mutation must first adapt to that mutation and survive to reproduce before any subsequent coevolution of the neural circuitry can occur. If ontogenetic connectivity depended on the phylogenetic *a priori ‘*knowledge’ embedded in chemically hardwired connectivity loops between muscles, that knowledge would be obsolete and potentially fatal with every new musculoskeletal mutation. In direct contrast, an evolutionary lineage that utilized a general solution to the control problem of novel biomechanics would gain a significant advantage for the evolution of new species.

## Authors contributions

JMDE, HJ and GEL designed the study and wrote the article. JMDE and HJ developed and tested the model of synaptic plasticity. AMJ, MK and JH worked on early versions of the musculoskeletal model.

